# Retinal ganglion cells undergo cell type–specific functional changes in a biophysically detailed model of retinal degeneration

**DOI:** 10.1101/2023.01.13.523982

**Authors:** Aiwen Xu, Michael Beyeler

## Abstract

Understanding the retina in health and disease is a key issue for neuroscience and neuroengineering applications such as retinal prostheses. During degeneration, the retinal network undergoes complex and multi-stage neuroanatomical alterations, which drastically impact the retinal ganglion cell (RGC) response and are of clinical importance. Here we present a biophysically detailed *in silico* model of retinal degeneration that simulates the network-level response to both light and electrical stimulation as a function of disease progression. The model is not only able to reproduce common findings about RGC activity in the degenerated retina, such as hyperactivity and increased electrical thresholds, but also offers testable predictions about the underlying neuroanatomical mechanisms. Overall, our findings demonstrate how biophysical changes associated with retinal degeneration affect retinal responses to both light and electrical stimulation, which may further our understanding of visual processing in the retina as well as inform the design and application of retinal prostheses.

## 1 Introduction

Understanding how the retina responds to light and electrical stimulation is a key issue for neuroscience and neuroengineering applications such as retinal prostheses. Computational models have been built either at the single-cell level or network level to understand the response properties of the healthy retina (for a recent review, see Guo et al. 2014). These include single-compartment models (“point models”) to simulate neuronal response as a function of ionic currents flowing across the neuronal membrane (e.g., Fohlmeister et al., 1990; Fohlmeister and Miller, 1997, Wohrer and Kornprobst, 2009), morphologically realistic models based on detailed anatomical representations of the physical components of biological neurons (e.g., Smith, 1995; Greenberg et al., 1999), and convolutional neural networks (e.g., McIntosh et al., 2016). Several studies did not just focus on the retina ‘s light response but also on the response to electrical stimulation (Cottaris and Elfar, 2005; Guo et al., 2016; Werginz et al., 2018; Beyeler, 2019; Paknahad et al., 2020), which may inform treatment options for people blinded by retinal degenerative diseases.

However, the retinal network undergoes drastic neuroanatomical alterations during retinal degeneration (Marc et al., 2003; Jones et al., 2016), which are of clinical importance to rehabilitative strategies such as retinal prostheses (Stingl et al., 2013; Luo et al., 2016; Palanker et al., 2020). These alterations are complex and multi-stage (Marc et al., 2003), starting with photoreceptor stress that leads to outer segment truncation (Phase I) and progressive photoreceptor cell death (Phase II), followed by a protracted period of global cell migration and cell death (Phase III). The consequences of these alterations on retinal ganglion cell (RGC) firing are manifold, which include hyperactivity (Pu et al., 2006; Stasheff, 2008; Stasheff et al., 2011; Telias et al., 2019), emergence of oscillations (Margolis et al., 2014; Ahn et al., 2022), and increased electrical stimulation thresholds (Rizzo et al., 2003b; O ‘Hearn et al., 2006; Jensen and Rizzo, 2008; Goo et al., 2011; Cho et al., 2016). With a few exceptions (Telias et al., 2019; Ahn et al., 2022), the underlying mechanistic reasons for most of these physiological changes remain poorly understood.

Previous computational work modeled retinal degeneration, but often stopped short of simulating the global retinal remodeling typified by the progressive nature of these diseases. For instance, Cottaris and Elfar (2005) built a model of the healthy retina and removed the cone population without addressing biophysical changes to the inner retina. Golden et al. (2018) simulated degeneration by removing a fraction of simulated neurons, increasing connectivity among the surviving neurons, and increasing the noise level, but did not address the progressive nature of these diseases. Other models stopped at reducing the thickness of different retinal layers (Paknahad et al., 2020, 2021) or hard-coded known physiological changes, such as increased spontaneous activity, into their model (Loizos et al., 2018). To the best of our knowledge, a comprehensive computational model of retinal degeneration is still lacking.

To address this, we built a biophysically inspired *in silico* computational model of the retina and simulated the network-level response to both light and electrical stimulation. After verifying that the model reproduced seminal findings about the light response of RGCs, we systematically introduced anatomical and neurophysiological changes to the network and studied their effect on network activity. In early phases of degeneration, we found that cone outer segment truncation and cone cell death differentially affected ON and OFF RGC firing: whereas the light response of ON RGCs diminished more quickly than that of OFF RGCs, the spontaneous firing rate of OFF RGCs steadily increased. In late phases of degeneration, we found that migratio and progressive death of inner retinal neurons led to a steady increase in electrical activation thresholds of both ON and OFF RGCs, especially for epiretinal stimulation. Our findings demonstrate how biophysical changes associated with retinal degeneration affect retinal responses to both light and electrical stimulation. A detailed model of the retina in health and disease has the potential to further our understanding of visual processing in the retina as well as inform the design of retinal prostheses.

## 2 Results

### 2.1. Light response of the healthy retina

Inspired by Cottaris and Elfar (2005), we started by simulating a three-dimensional healthy patch (300 µm × 300 µm × 210 µm) of the parafoveal retina (see Methods). The network consisted of 11, 138 cells belonging to nine different cell types (4, 149 cones, 537 horizontal cells, 3, 508 ON/OFF bipolar cells, 779 ON/OFF wide-field amacrine cells, 723 narrow-field amacrine cells, and 1, 442 ganglion cells), connected via generally accepted (Rodieck, 1998; Wassle and Boycott, 1991) synaptic connections (see Fig. 1).

**Figure 1.**
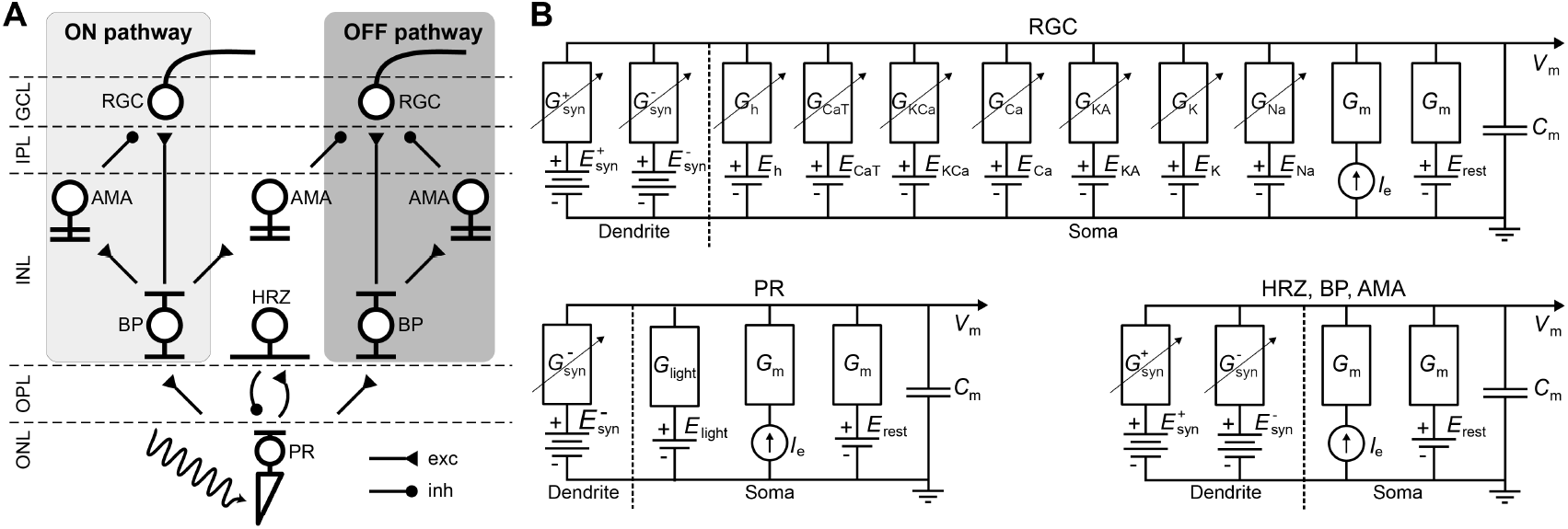
**A)** Diagram of the connections between the retinal neurons in the healthy state. PR: photoreceptor, HRZ: horizontal cell, BP: bipolar cell, AMA: amacrine cell, RGC: retinal ganglion cell. ONL: outer nuclear layer, OPL: outer plexiform layer, INL: inner nuclear layer, IPL: inner plexiform layer, GCL: ganglion cell layer. **B)** RC circuit model of a neuron ‘s membrane potential. All neurons included a membrane capacitance (*C*_*m*_), a leakage current (*I*_leak_), an external current driven by the extracellular potential gradient (*I_e_*), and synaptically gated ionic currents (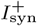 and 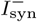). RGCs had additional voltage-gated and ligand-gated ionic currents (Gu et al., 2016) and photoreceptors had a photo-sensitive current (*I*_light_) as described in Cottaris and Elfar (2005). Dendritic trees and ganglion cell axons were not modeled.

Briefly, cone photoreceptors (labeled “PR” in Fig. 1A) excited ON and inhibited OFF bipolar cells (“BP”). In addition, they excited horizontal cells (“HRZ”), which in turn inhibited cone terminals, thus generating an inhibitory surround in the bipolar cell response. ON and OFF bipolar cells then excited ON and OFF amacrine cells (“AMA”) as well as ON and OFF ganglion cells (“RGC”), respectively, to generate an inhibitory surround in the ganglion cell response. ON and OFF amacrine cells also provided lateral inhibition to ON and OFF ganglion cells, respectively. Lastly, we included a unilateral inhibitory connection from a special type of narrow-field ON amacrine cell to OFF ganglion cells (Wyatt and Rizzo, 1996). Rod circuitry was not implemented.

To implement the biophysical properties of retinal neurons, we largely followed Cottaris and Elfar (2005) to modify a leaky integrator model by adding membrane and synaptic conductances (Fig. 1B). We assumed that neurons are electronically compact, so their activation levels could be described by a single membrane potential. All 11, 138 neurons had a spatially nonzero soma with non-gated ion channels (leakage channels) modeled by a constant linear conductance (*G*_m_) in series with a constant single-cell battery (i.e., the cell ‘s resting voltage, *E*_rest_) and an extracellular current that modeled extracellular electrical stimulation (Eq. 1). Synapses were assumed to lie at the center of a neuron ‘s dendritic field (excitatory if *E*_syn_ *> E*_rest_ and inhibitory if *E*_syn_ < *E*_rest_). In addition, cones had a photosensitive ion channel (Eq. 3). Ganglion cells deviated from Cottaris and Elfar (2005) and were implemented as Hodgkin-Huxley neurons with seven ion channels (Eq. 6) that were previously shown to capture the firing dynamics of RGCs in the rabbit retina (Guo et al., 2016).

Neurons were arranged in nine stacked hexagonal mosaics (Fig. 2A). Here the retina reacted to a bright disk stimulus (40 µm in radius, with light intensity 1.0) presented against a gray background (300 µm by 300 µm, with light intensity 0.5). Fig. 2B breaks down the retinal response to the disk stimulus layer by layer. Upon stimulus onset, photoreceptors in the center of the mosaic became hyperpolarized and were surrounded by a thin ring of depolarized cells due to reduced lateral inhibition provided by the horizontal cells. This activity spread through both ON and OFF pathways, leading to depolarized ON bipolar cells and spiking ON RGCs, whereas corresponding cells in the OFF pathway were hyperpolarized.

**Figure 2.**
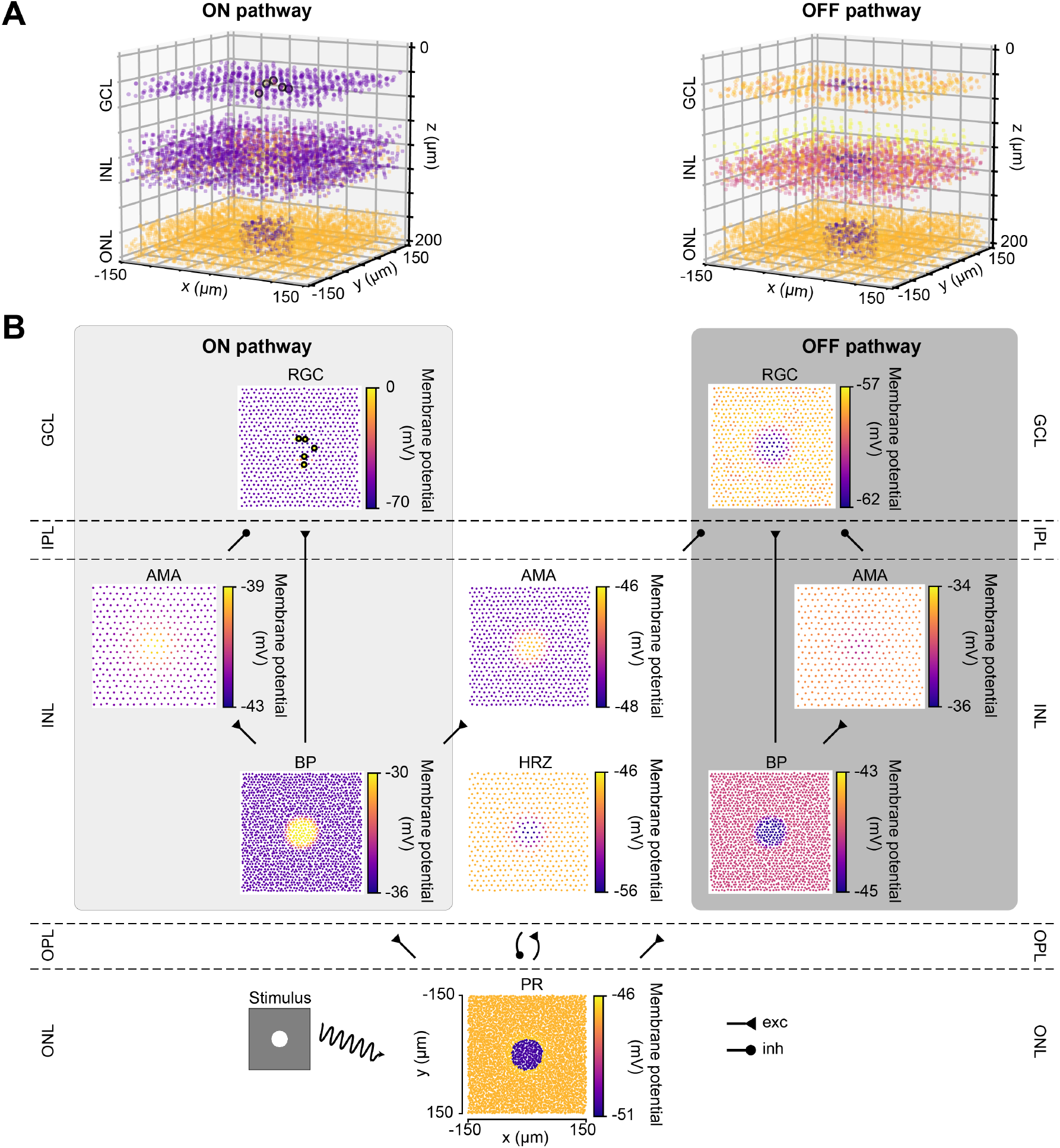
The light response of the retinal network in the healthy state, presented both in 3D and by cell type. Abbreviations same as in Fig. 1. **A)** The light response of the healthy retina presented separately for the ON pathway (left) and the OFF pathway (right) in 3D. The light stimulus that was used to elicit the response was a bright disk (40 *µ*m in radius with light intensity 1) placed at the center of a gray background (300 *µ*m by 300 *µ*m with light intensity 0.5) (illustrated by the bottom left inset of *B*). The light response shown occured 110 ms after stimulus onset. Each circle represents the (*x, y, z*) location of a neuron, and the color of each circle indicates the membrane potential of each neuron, with the color bars in *B*. Enlarged circles with black border indicate neuronal spikes. The plot of the ON pathway includes cone photoreceptors, horizontal cells and all the cells belonging to the ON pathway, and the plot of the OFF pathway includes cone photoreceptors, horizontal cells and all the cells belonging to the OFF pathway. **B)** The light response presented by cell type, corresponding to each layer in *A*. Each circle represents the x-y location of a neuron, and the color of each circle indicates the membrane potential of each neuron. Enlarged circles with black border indicate neuronal spikes. For an animated version of this figure, see Supplemental Video 1.

The spatiotemporal evolution of RGC activity in response to the above mentioned stimulus is shown in Fig. 3, which was modeled after Fig. 5 in Cottaris and Elfar (2005). The stimulus mentioned above was modulated in time by a square wave signal (200 ms phase duration) at four contrast levels: 100 %, 50 %, −50 %, −100 %. Consistent with conduction delays in the rabbit retina (Zeck et al., 2011), ON RGCs first fired roughly 20 ms after stimulus onset, whereas OFF RGCs took 50 ms to respond (Fig. 3A–B). RGC unaffected by the stimulus exhibited at a 2 Hz spontaneous firing rate, which was achieved by setting the conductance of the leakage current (see Section 4.1). Synchronization of firing increased with stimulus strength for both ON and OFF populations. The center-surround structure of RGC receptive fields is evident in Fig. 3C–D. ×

**Figure 3.**
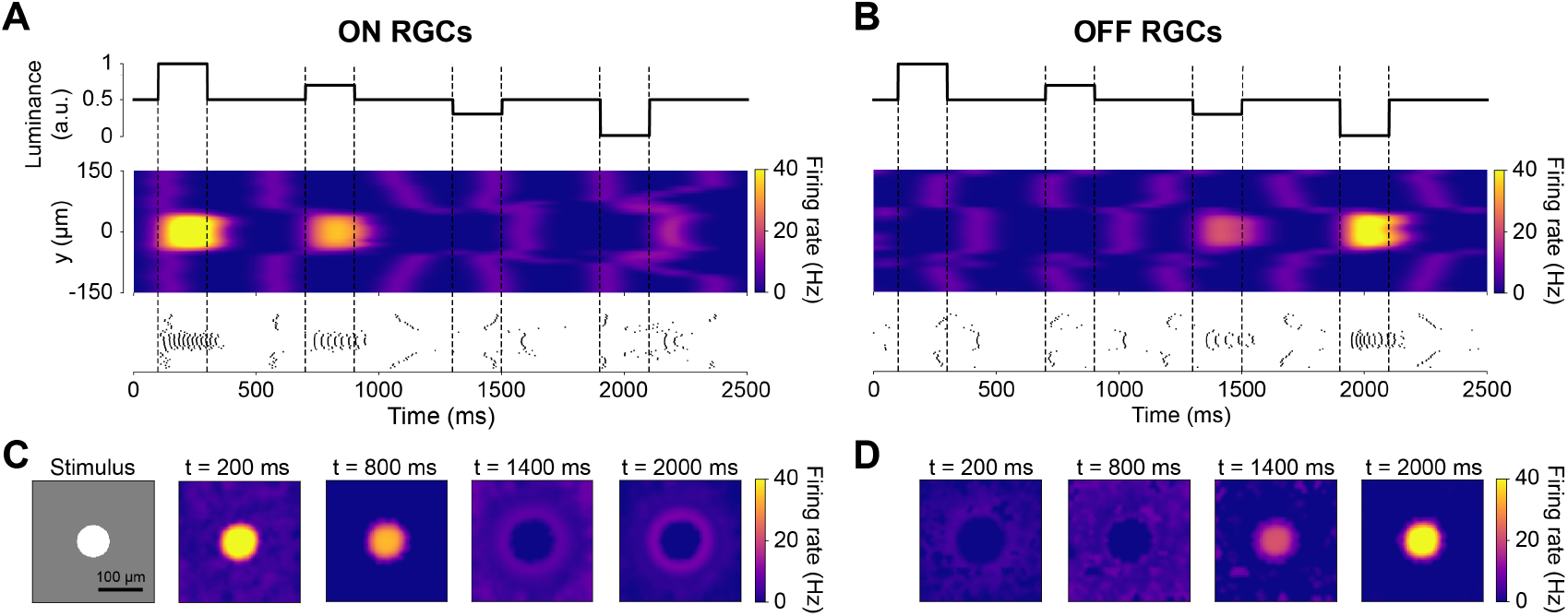
The spatiotemporal response of RGCs to a temporally varying light stimulus (inspired by Cottaris and Elfar, 2005). The light stimulus (illustrated in the bottom-left inset) was a disk (40 µm in radius) placed at the center of a gray background (300 µm × 300 µm with light intensity 0.5), varying in intensity over time. **A–B)** Spatiotemporal profile of RGC firing rate for neurons located at *x* = 0 µm, visualized both as a heatmap (smoothed with a 50 ms Gaussian sliding window) and raster plot. The vertical dotted lines indicate the time of change in light intensity of the disk stimulus. **C–D)** Spatial activity profiles of RGC firing rate taken at different points in time.

### 2.2. Retinal degeneration differentially affects the spontaneous firing of ON and OFF cells

After verifying the light response of the healthy retina model, we gradually introduced neuroanatomical and neurophysiological changes to the network in order to model retinal degeneration (Fig. 4). Retinal degenerative diseases are commonly described in the literature as progressing in three phases (Marc et al., 2003; Jones et al., 2016), starting with photoreceptor stress that leads to outer segment truncation (Phase I) and progressive photoreceptor cell death (Phase II), followed by a protracted period of global cell migration and cell death (Phase III). To make the modeling of such a complex process feasible, we limited ourselves to the major changes that may have an impact on RGC signaling as outlined below (see Section 4.3 for details). Because our model did not include rods, we combined Phases I and II into Phase I/II, where we gradually reduced the cone population and shortened the outer segments of the surviving cones (Fig. 4A). The complete loss of photoreceptors and horizontal cells marked the end of Phase I/II (Fig. 4B). To simulate global retinal remodeling in Phase III, we gradually reduced the population of bipolar and amacrine cells while migrating a randomly chosen fraction of inner retinal neurons to different retinal layers (Fig. 4C). To model disease progression over time (Fig. 4E), we assumed a linear reduction in cone segment length and cone population during Phase I/II, and a linear reduction in cell survival rate as well as a linear increase in cell migration rate during Phase III.

**Figure 4.**
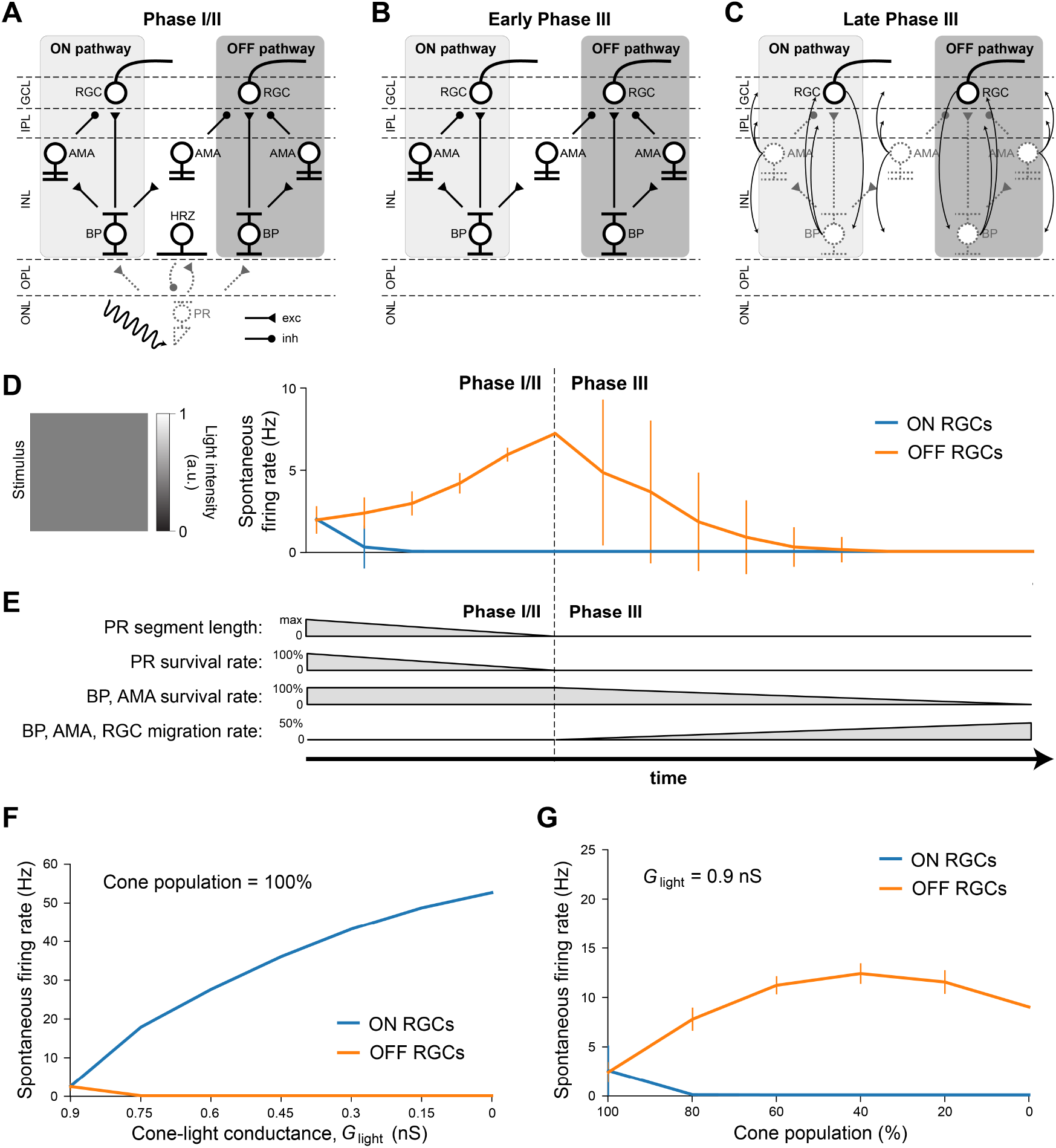
Simulating retinal degeneration. Abbreviations same as in Fig. 1. **A)** To simulate Phase I/II of retinal degeneration, we gradually shortened the cone outer segment length while simultaneously reducing the cone population. **B)** The complete loss of the cone population marks the beginning of Phase III. **C)** During Phase III, we gradually reduced the population of bipolar and amacrine cells. In addition, a fraction of surviving cells migrated to different layers: amacrine cells started to migrate to the horizontal cell layer, the inner plexiform layer, and the ganglion cell layer; bipolar cells started to migrate to the inner plexiform layer and the ganglion cell layer; RGCs started to migrate to the horizontal cell layer. At the end of Phase III, all inner retinal neurons have degenerated. **D)** Spontaneous firing rate of ON and OFF RGCs as a function of disease progression. Input was a full-field stimulus of *L*(*t*) = 0.5 light intensity (Eq. 5). Values indicate the spontaneous firing rate measured over 2500 milliseconds and averaged across all RGCs, with vertical bars indicating the standard deviation. **E)** To simulate disease progression over time, we assumed a constant rate of change for PR segment length and neuron survival. **F)** RGC spontaneous firing rate as a function of cone outer segment truncation (simulated as a reduction in the cone-light conductance *G*_light_, see Eq. 5), averaged across RGCs, while the cone population size was held constant. **G)** RGC spontaneous firing rate as a function of the size of the cone population, averaged across the population of surviving RGCs, while *G*_light_ was held constant.

**Figure 5.**
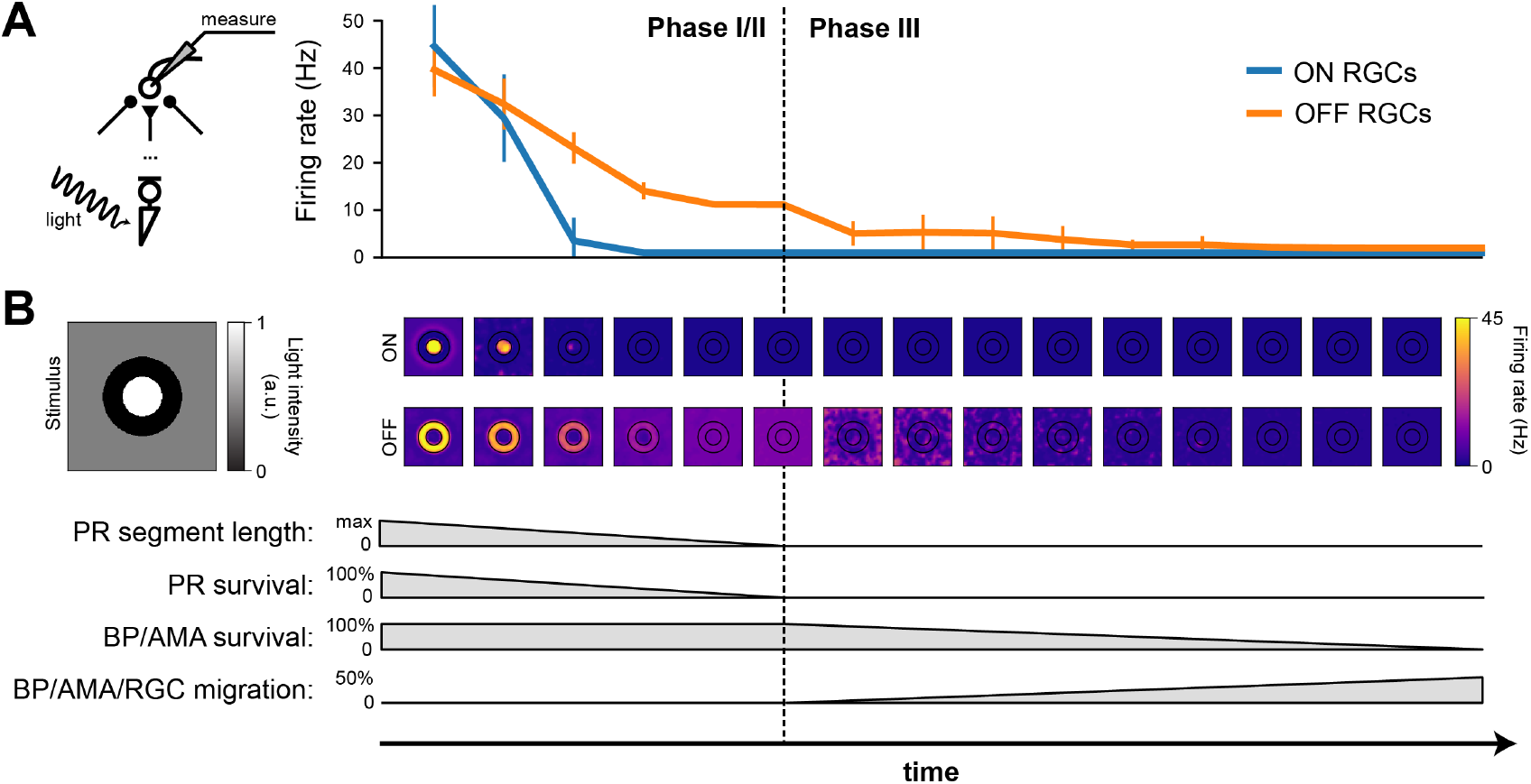
RGC firing rate (in response to light stimulation) decreases with the progression of degeneration. The stimulus was a bright disk (40 µm in radius with light intensity 1) surrounded by a dark ring (40 µm in inner radius and 80 µm in outer radius with light intensity 0) placed on a gray background (300 µm by 300 µm with light intensity 0.5). **A)** Firing rate of ON and OFF RGCs, measured over the 1000 ms stimulus presentation, plotted against time. The mean and standard deviation were calculated from the ON RGCs under the bright disk and the OFF RGCs under the dark ring. **B)** The spatial activity of ON RGCs (top row) and OFF RGCs (bottom row) plotted at different time points of degeneration. The borders of the bright center and dark surround are outlined in black.

Although we did not alter the inherent excitability of RGCs, the network-level changes described above had a profound impact on RGC activity. Fig. 4D shows the spontaneous firing rate of RGCs as a function of disease progression, measured over 2500 milliseconds and averaged across all RGCs. Whereas most ON RGCs were silenced as soon as the cone population and outer segment length dropped below 80 % of their initial values, OFF RGCs experienced a gradual increase in spontaneous firing rate throughout Phase I/II, which peaked at roughly 300 % of its initial value at the beginning of Phase III. After that, the average spontaneous firing rate of the OFF RGC population gradually dropped to zero, although individual cells differed greatly in their activation profiles. Despite the increased variability in mean activity, no bursting or oscillatory activity emerged, as evidenced by Poissonian inter-spike interval distributions (Fig. A1).

We identified the underlying mechanistic causes of the change in spontaneous firing rate and found that they differed for ON and OFF RGCs. Whereas ON cells increased their spontaneous firing mainly as a function of cone outer segment truncation (Fig. 4F), OFF cells were mainly affected by the size of the surviving cone population (Fig. 4G). The full range of light responses as a function of outer segment truncation and cone population size is given in Fig. A2.

### 2.3. Light response of ON cells decreases more quickly than that of OFF cells during degeneration

To investigate how the light response of RGCs changed as a function of disease progression, we presented a constant stimulus in all stages of the disease (Fig. 5). The stimulus was a bright disk (40 µm in radius with maximal light intensity, *l*(*t*) = 1, Eq. 5) surrounded by a dark ring (40 µm in inner radius and 80 µm in outer radius with light intensity 0) placed on a gray background simulated with light intensity 0.5 (Fig. 5B, *left*). The stimulus was presented for 1000 milliseconds, during which the mean firing rate of each RGC was calculated.

Whereas both ON and OFF cells initially responded with similar firing rates, ON cells saw a much quicker reduction in firing rate during Phase I/II than OFF cells, remaining silent for the second half of Phase I/II and all throughout Phase III (Fig. 5A).

The spatial response profile at different time steps (where the center of the image is aligned with the corresponding time of disease progression on the *x* axis) is shown Fig. 5B. Whereas the center-surround structure of the retinal response is preserved during early stages of Phase I/II, spatial specificity is quickly lost during later stages of Phase I/II. During Phase III, it is not uncommon for the most active OFF cells to be found far away from the site of stimulation.

### 2.4. Ganglion cells quickly lose spatial selectivity during Phase I/II

To further illustrate the change in spatiotemporal receptive field (RF) profiles, we fit a generalized linear model (GLM) to the spiking response of RGCs to a “cloud” stimulus consisting of spatiotemporal Gaussian white noise filtered with a two-dimensional spatial Gaussian filter (Shi et al., 2019; for details see Section 4.4).

The fitted spatiotemporal RFs of two example cells located at the center of the simulated retinal patch is shown in Fig. 6. As expected, the RF profile of the healthy ON cell showed a clear excitatory center and an inhibitory surround at the time of a spike (*t* = 0 ms), with reversed polarity at *t* = −100 ms. In early Phase I/II (where cone population and outer segment length were at 80 % of their healthy values), inhibitory subregions at times *t <* 0 were much broader and much more pronounced, and were followed by an enlarged excitatory center that had lost its circular shape. In later stages of Phase I/II (where cone population and outer segment length were at 40 % of their healthy values), the ON cell was no longer sufficiently responsive to light stimulationn and its spatial response profile was lost.

**Figure 6.**
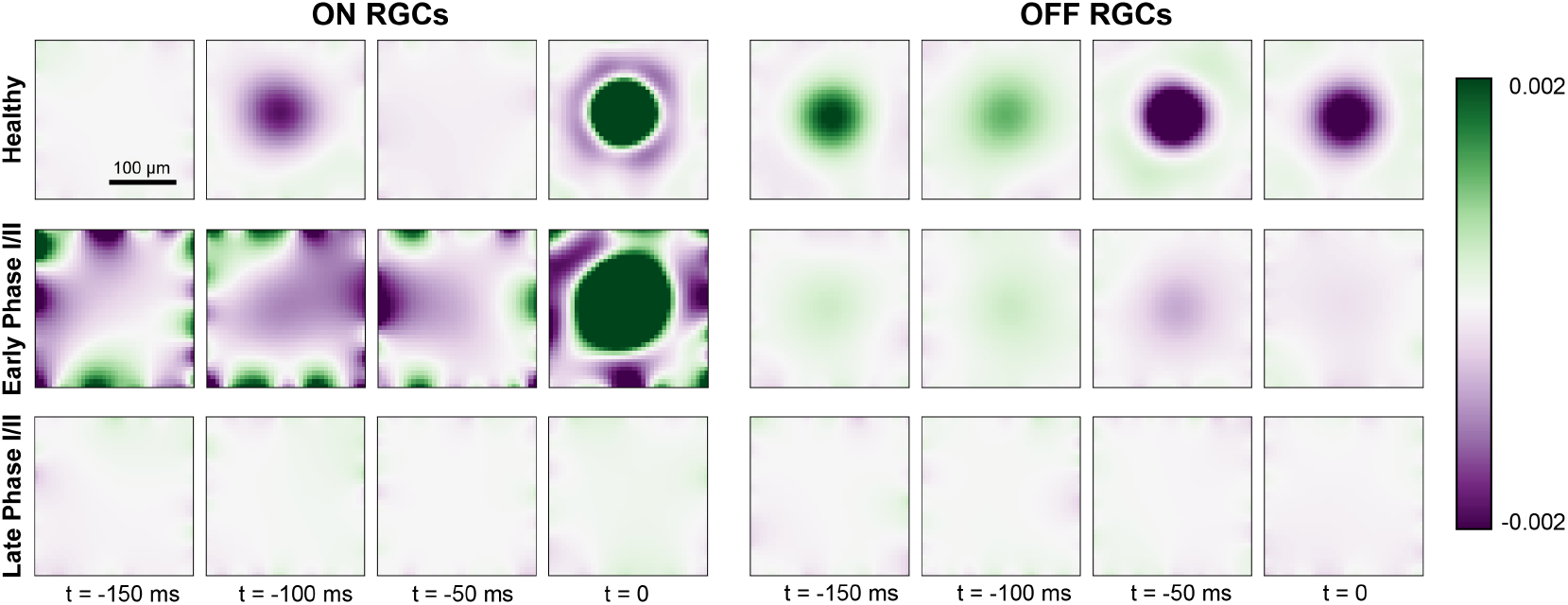
GLM fit for ON cells (*left*) and OFF cells (*right*) for a healthy retina (*top row*), early Phase I/II (*middle row* ; 100 % cones surviving and *G*_light_ = 0.75 for ON, 60 % cones surviving and *G*_light_ = 0.6 for OFF), and late Phase I/II (*bottom row* ; 80 % cones surviving and *G*_light_ = 0.75 for ON, 40 % cones surviving and *G*_light_ = 0.45). The colorbar represents the range of values of the linear filters *k_i_* (see 4.4) at each spatial and temporal location, where green indicates excitatory values and purple indicates inhibitory values.

The RF profile of the OFF cell also exhibited a center-surround structure, though the excitatory surround was most pronounced at *t* = −50 ms due to the longer synaptic delay of the OFF pathway (see Section 4.2). This also led to a prolonged response at the center of the RF profile. In early Phase I/II, the spatial response profile of the OFF cell seemed to be largely preserved, although the overall response as weakened. In late Phase I/II, the OFF cell was no longer sufficiently responsive to light stimulation and its spatial response profile was lost.

### 2.5. Electrical thresholds increase throughout retinal degeneration

Understanding how the degenerated retina responds to electrical stimulation is crucial for treatment options such as retinal prostheses. We thus placed a simulated disk electrode (80 µm) either epiretinally (i.e., 2 µm above the ganglion cell layer; Cottaris and Elfar 2005) or subretinally (i.e., at *z* = 135 µm, close to horizontal and bipolar cells) and measured the RGC response to a 20 Hz cathodic-first biphasic pulse train of 1 s duration with 0.45 ms phase duration (Fig. 7).

**Figure 7.**
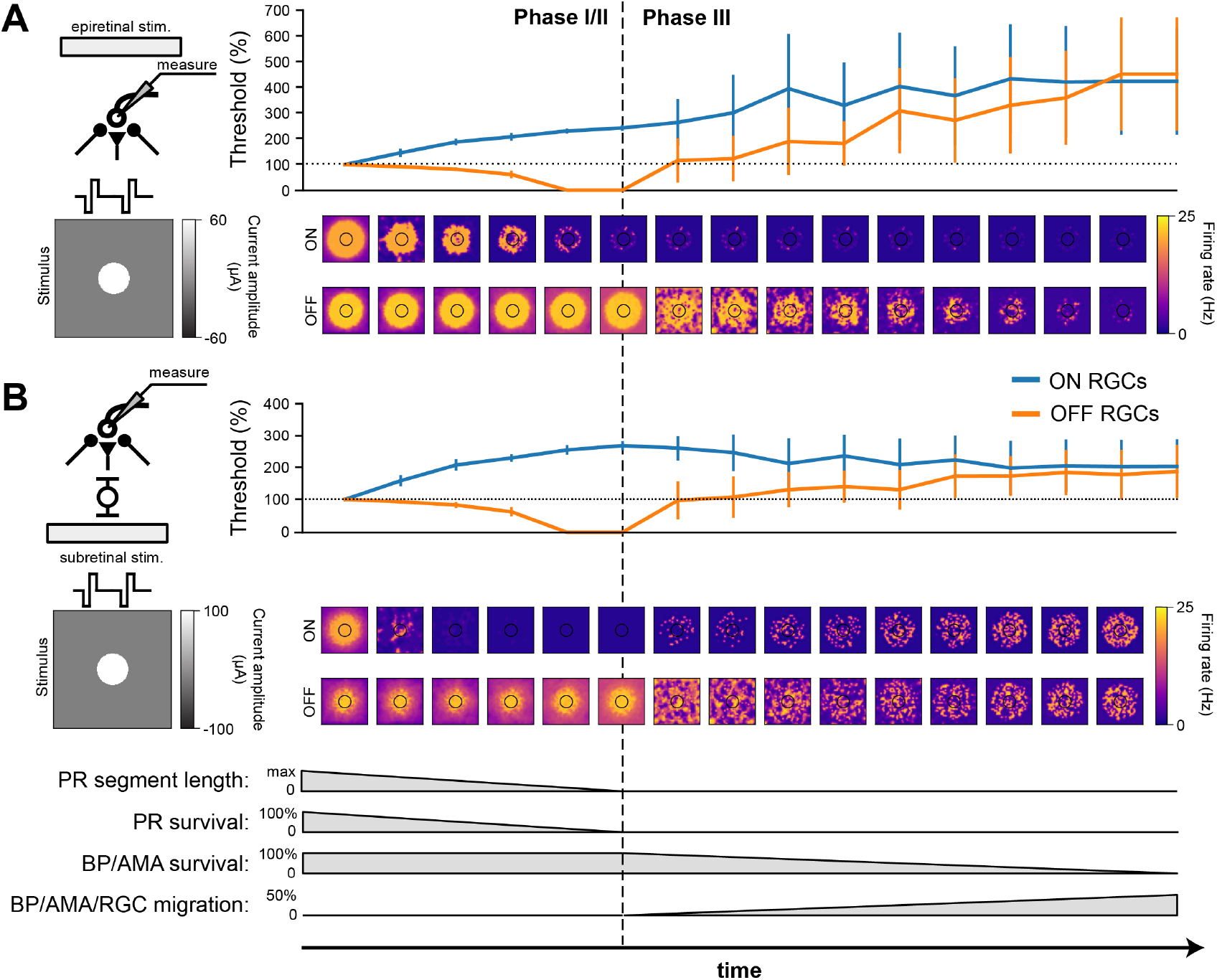
RGC response to electrical stimulation with a 20 Hz biphasic cathodic-pulse train (0.45 ms phase duration, 1 s stimulus duration). Mean values were calculated over 1000 ms (stimulus presentation) and averaged across neurons located directly below the electrode; vertical bars the SD. **A)** Epiretinal stimulation. Electrical stimulation thresholds during retinal degeneration reported relative to healthy thresholds (100 %, horizontal dotted line). The spatial response profile of ON and OFF RGCs is given below for a pulse train of 60 µA amplitude. The borders of the electrode are outlined in black. **B)** Subretinal stimulation. Electrical stimulation thresholds during retinal degeneration reported relative to healthy thresholds (100 %, horizontal dotted line). The spatial response profile of ON and OFF RGCs is given below for a pulse train of 60 µA amplitude. The borders of the electrode are outlined in black.

Consistent with the literature (Rizzo et al., 2003b; O ‘Hearn et al., 2006; Jensen and Rizzo, 2008; Goo et al., 2011; Cho et al., 2016), electrical thresholds in later stages of degeneration rose to 200 %–400 % of those in the healthy retina (Fig. 7). Here, threshold was defined as the smallest stimulus amplitude that elicited a spike on at least half of 20 trials (Sekirnjak et al., 2009; Weitz et al., 2014), and the resulting thresholds were averaged across the population of surviving RGCs.

Interestingly, our model predicted that degeneration should affect the electrical thresholds of ON and OFF RGCs differently: whereas thresholds for ON cells tended to rise rapidly in Phase I/II, OFF cell thresholds decreased throughout Phase I/II to a point where the threshold was effectively zero, due to increased spontaneous firing. As cells started to migrate in Phase III, thresholds rose again. Notably, epiretinal stimulation thresholds kept rising (Fig. 7A, reaching thresholds up to 400 % of those found in a healthy retina, whereas subretinal thresholds were more stable during Phase III and plateaued at around 200 % of the healthy thresholds (Fig. 7). The standard deviations in Phase III were significantly larger than those in Phase I/II because of the cell migration in Phase III.

In addition, the spatial response profiles were strongly affected by degeneration (heat maps in Fig. 7, here shown for a 20 Hz biphasic pulse train with 60 mA amplitude). A ring-like structure is noticeable in both ON and OFF cell responses due to the influence of the extracellular potential being highest along the edge of the electrode, allowing activity to spread far beyond the size of the electrode. Epiretinal stimulation was able to elicit solid OFF cell responses throughout Phase I/II, whereas ON cell responses lasted only halfway through. After that, cell migration started to disrupt spatial response profiles in Phase III. The story was similar for subretinal stimulation, though a few differences were noticeable. First, ON cell responses vanished almost immediately in Phase I/II, only to come back in late stages of Phase III as more and more ganglion cells started to migrate closer to the subretinal electrode. Second, the spatial activation profile of OFF cells was more confined early in Phase I/II, but began to widen due to increased spontaneous RGC later in Phase I/II. Third, cell migration disrupted the spatial response profiles more quickly and more thoroughly, but continued to elicit responses all the way to the end of Phase III.

### 2.6. Cell death and migration affect ON and OFF cells differently

To isolate the network changes responsible for the altered electrical response properties of RGCs, we simulated frequency-current (F-I) curves at different stages of degeneration for three different stimulation modes: current injection, epiretinal electrical stimulation, and subretinal electrical stimulation (Fig. 8). During Phase I/II, cone death reduced the response of ON cells for all three stimulation modes (Fig. 8, *top row*), but left OFF cells unaffected. This is consistent with the retina ‘s light response in Fig. 4F. During Phase III, bipolar and amacrine cell death reduced the response of OFF cells for all three stimulation modes, but left the ON cells mostly unaffected (Fig. 8, *top row*); the one exception being subretinally stimulated ON cells, which saw the greatest reduction in activity. Finally, cell migration affected the response of both ON and OFF cells for both epiretinal and subretinal stimulation, increasing response variability across the RGC population.

**Figure 8.**
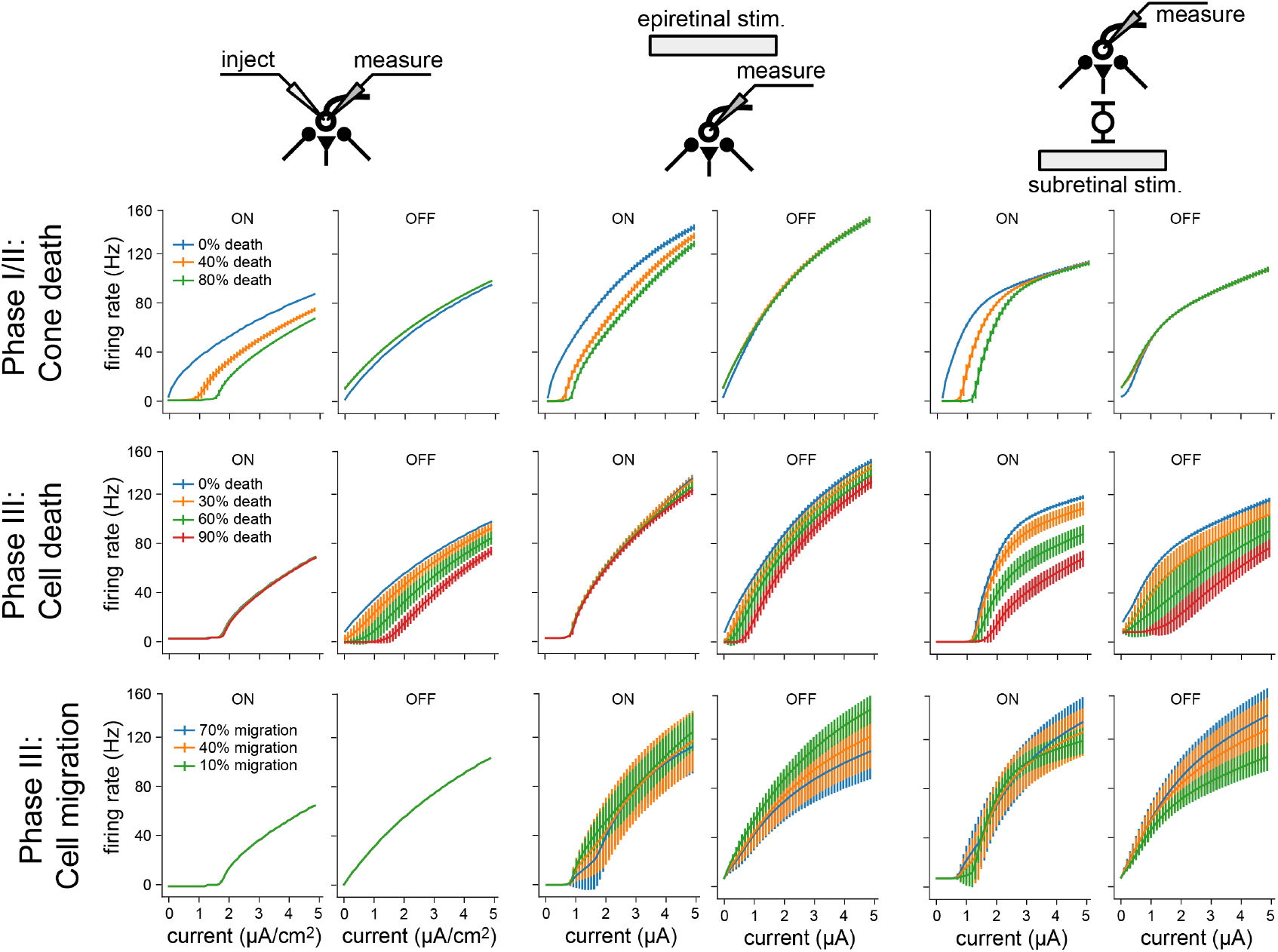
Frequency-current (F-I) curves for three modes of neuronal stimulation: current injection (*two leftmost columns*), epiretinal electrical stimulation (*two center columns*) and subretinal electrical stimulation (*two rightmost columns*). F-I curves are shown for RGCs at different stages of retinal degeneration: as a function of cone death during Phase I/II (*top row*), as a function of bipolar and amacrine cell death during Phase III (*middle row*), and as a function of bipolar, amacrine, and ganglion cell migration during Phase III (*bottom row*). Values averaged across RGCs; vertical bars are the standard deviation.

Overall, these results demonstrate how network-level changes may affect RGC firing at different stages of retinal degeneration.

## 3 Discussion

We have developed a biophysically inspired *in silico* model of retinal degeneration that simulates the network-level response to both light and electrical stimulation, and found that degeneration differentially affects ON and OFF RGCs. Existing computational models of retinal degeneration largely focus on RGC activity in the absence of photoreceptor input (e.g., Cottaris and Elfar, 2005; Golden et al., 2018), but do not consider the global retinal remodeling that may impact the responsiveness of RGCs. To this end, our simulations do not just reproduce commonly reported findings about the changes in RGC activity encountered during retinal degeneration (e.g., hyperactivity, increased electrical thresholds) but also offer testable predictions about the neuroanatomical mechanisms that may underlie altered RGC activity as a function of disease progression.

Consistent with the literature (Pu et al., 2006; Stasheff, 2008; Sekirnjak et al., 2011; Stasheff et al., 2011; Telias et al., 2019), we found that RGCs exhibited elevated spontaneous firing rates during retinal degeneration (Fig. 4). In our model this was mainly restricted to the OFF RGC population, which became more active over time (Fig. 4D, G). Similar observations have been made in degenerated retinas of mouse models (Pu et al., 2006; Stasheff, 2008; Sekirnjak et al., 2011), which are dominated by OFF cell activity. However, we identified photoreceptor cell death in Phase I/II as the main driving force behind this hyperactivity, as opposed to an intrinsic change to RGC excitability (Telias et al., 2019). Moreover, the complete loss of cones did not drive RGCs into an oscillatory state (Fig. A2). On the other hand, ON cells showed increased activity in response to cone outer segment truncation (Fig. 4F), which may occur only early in degeneration before most photoreceptors are lost, after which their spontaneous activity is expected to quickly vanish.

As the light response of the RGC population slowly subsided (Fig. 5), ON cells saw a much quicker reduction in firing rate than OFF cells, remaining silent for the second half of Phase I/II and all throughout Phase III (Fig. 5). GLM fits showed a brief broadening of the spatiotemporal RF for ON RGCs early in Phase I/II while losing their inhibitory surround, before the spatial properties of both ON and OFF RFs were lost (Fig. 6). This is consistent with studies that have documented cell type–specific functional changes in RGCs across animal models (Pu et al., 2006; Sekirnjak et al., 2011; Yu et al., 2017), where spatial receptive fields often lose their circular shape and appear spotty before they vanish (Yu et al., 2017).

As degeneration progressed, electrical thresholds tended to increase for both subretinal and epiretinal stimulation (Fig. 8), which is consistent with most literature on the subject (Rizzo et al., 2003b; O ‘Hearn et al., 2006; Jensen and Rizzo, 2008; Goo et al., 2011; Cho et al., 2016)—though see Sekirnjak et al. (2009). Mirroring the changes in the light response, ON cells also displayed diminished responses to electrical stimulation (Fig. 8). This resulted in higher activation thresholds for ON cells as compared to OFF cells (Fig. 7), which is a phenomenon previously documented in the degenerated mouse retina (Yang et al., 2018). Interestingly, our model also predicts a brief period of degeneration during which OFF cells are so active that their electrical threshold is effectively zero. Furthermore, our model offers testable predictions of how cone death (Phase I/II), bipolar and amacrine cell death (Phase III), and cell migration (Phase III) affect the responsiveness of RGCs (Fig. 8).

Overall, our findings demonstrate how biophysical changes associated with retinal degeneration affect retinal responses to both light and electrical stimulation, which may have important implications for the design and application of retinal prostheses.In specific, our results suggest that spatially confined responses might be more easily elicited with subretinal stimulation in Phase I/II and with epiretinal stimulation in Phase III (Fig. 7), though cell migration might strongly affect both stimulation modes (Fig. 8). Moreover, both subretinal and epiretinal stimulation are expected to more easily activate OFF cells, as they remain active through most of Phase III. This suggests that OFF cell activity may play a greater role in prosthetic vision than previously assumed (Im and Fried, 2015).

Although our model captures a range of biophysical changes common to retinal degeneration, the complex nature of this process necessitated some simplifying assumptions. For instance, we only implicitly modeled the effect of Muller cell hypertrophy on RGC activity and did not attempt to model the formation of microneuromas and other morphological changes (Marc et al., 2003). In addition, as we did not modify the intrinsic properties of the RGC population, our results suggest that commonly documented physiological changes such as RGC hyperactivity and increased electrical thresholds (Chen et al., 2013; Saha et al., 2016; Telias et al., 2019) may have additional network-mediate causes that are presynaptic to RGCs. This work thus offers testable predictions to further our understanding of retinal processing in health and disease.

## 4. Methods

The retina model was implemented using Brian 2 (Stimberg et al., 2019) and Brian2GeNN (Stimberg et al., 2020) in Python. All simulations were run on a single NVIDIA RTX 3090 (24 GB of GPU memory). All our code will be made publicly available once the manuscript is accepted.

### 4.1. Modeling individual neurons

The healthy retinal network consisted of neurons belonging to nine different cell types: cones, horizontal cells, ON/OFF bipolar cells, ON/OFF wide-field amacrine cells, narrow-field amacrine cells, and ON/OFF RGCs. All neuronal parameters can be found in Tables 1 and 2. Synaptic parameters are given in Table 3. In the below equations, variables are generally denoted by lowercase letters and constants are denoted by uppercase letters.

**Table 1.** Neuronal membrane parameters. *G*_m_ for RGCs was set such that the spontaneous firing rate was around 2 Hz. All other values for *G*_m_ and *C*_m_ were adopted from Cottaris and Elfar (2005). All others were adopted from Guo et al. (2016).

|  | PR | HRZ | BP<sub>ON/OFF</sub> | AMA<sub>ON/OFF</sub><sup>WF/NF</sup> | RGC<sub>ON</sub> | RGC<sub>OFF</sub> |
| --- | --- | --- | --- | --- | --- | --- |
| $C_{\text{m}}$ (pF) | 80.0 | 210.0 | 50.0 | 50.0 | 50.0 | 50.0 |
| $G_{\text{m}}$ (nS) | 4.0 | 2.5 | 2.0 | 2.0 | — | — |
| $G_{\text{m}}$ (mS cm<sup>-2</sup>) | — | — | — | — | 0.3 | 0.274 |
| $G_{\text{light}}$ (nS) | 0.9 | — | — | — | — | — |
| $E_{\text{rest}}$ (mV) | -50.0 | -65.0 | -45.0 | -50.0 | -66.5 | -70.5 |
| $E_{\text{Na}}$ (mV) | — | — | — | — | 35.0 | 35.0 |
| $E_{\text{K}}$ (mV) | — | — | — | — | -72.0 | -68.0 |
| $E_{\text{h}}$ (mV) | — | — | — | — | -45.8 | -26.8 |
| $G_{\text{Na}}$ (mS cm<sup>-2</sup>) | — | — | — | — | 1072.0 | 249.0 |
| $G_{\text{K}}$ (mS cm<sup>-2</sup>) | — | — | — | — | 40.5 | 68.85 |
| $G_{\text{KA}}$ (mS cm<sup>-2</sup>) | — | — | — | — | 94.5 | 18.9 |
| $G_{\text{Ca}}$ (mS cm<sup>-2</sup>) | — | — | — | — | 2.1 | 1.6 |
| $G_{\text{KCa}}$ (mS cm<sup>-2</sup>) | — | — | — | — | 0.04 | 0.0474 |
| $G_{\text{h}}$ (mS cm<sup>-2</sup>) | — | — | — | — | 0.4287 | 0.1429 |
| $G_{\text{CaT}}$ (mS cm<sup>-2</sup>) | — | — | — | — | 0.008 | 0.1983 |

**Table 2.**
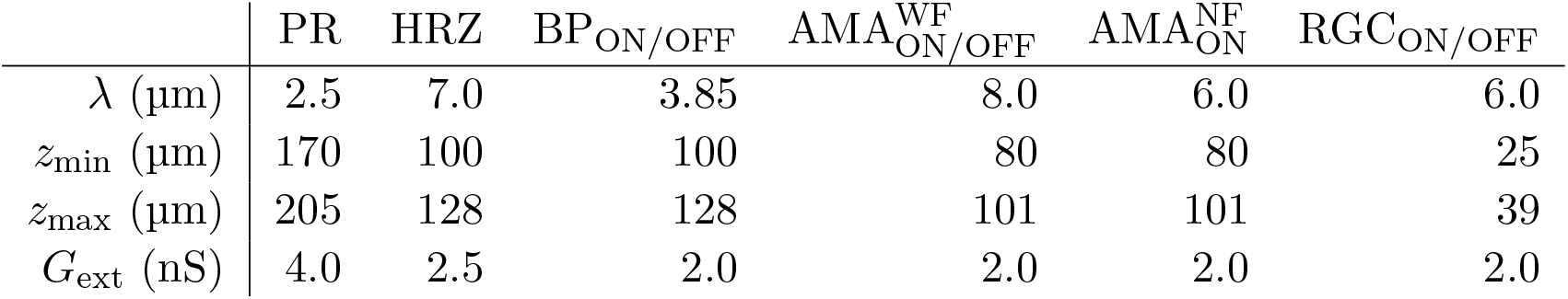
Spatial layout parameters for the different cell types (Cottaris and Elfar, 2005). PR: photoreceptor, HRZ: horizontal cell, BP: bipolar cell, AMA: amacrine cell, RGC: retinal ganglion cell, WF: widefield, NF: nonwide-field.

**Table 3.** Synaptic parameters. The synaptic delays were set such that the latency of ON RGCs were around 20 ms and the latency of OFF RGCs were around 50 ms. All other values are from Cottaris and Elfar (2005).

| | $\tau$ (ms) | $E_{\text{syn}}$ (mV) | $G_{\text{min}}$ (nS) | $G_{\text{max}}$ (nS) | $V_{50}$ (mV) | $\beta$ (mV) | type | $\sigma$ ($\mu\text{m}$) |
| --- | --- | --- | --- | --- | --- | --- | --- | --- |
| PR→HRZ | 7 | 0.0 | 0.0 | 7.0 | -43.0 | 2.0 | I | 10.5 |
| HRZ→PR | 7 | -67.0 | 0.0 | 3.0 | -29.5 | 7.4 | I | 2.5 |
| PR→BP<sub>ON</sub> | 5 | 0.0 | 0.1 | 1.1 | -47.0 | 1.7 | D | 3.85 |
| PR→BP<sub>OFF</sub> | 13 | 0.0 | 0.0 | 3.75 | -41.5 | 1.2 | I | 3.85 |
| BP<sub>ON</sub>→AMA<sub>ON</sub><sup>WF</sup> | 5 | 0.0 | 0.0 | 1.0 | -33.5 | 3.0 | I | 24.0 |
| BP<sub>ON</sub>→AMA<sub>ON</sub><sup>NF</sup> | 5 | 0.0 | 0.0 | 0.2 | -35.0 | 3.0 | I | 6.0 |
| BP<sub>OFF</sub>→AMA<sub>OFF</sub><sup>WF</sup> | 11 | 0.0 | 0.0 | 1.8 | -44.0 | 3.0 | I | 24.0 |
| BP<sub>ON</sub>→RGC<sub>ON</sub> | 5 | 0.0 | 0.0 | 2.5 | -33.5 | 3.0 | I | 6.0 |
| AMA<sub>ON</sub><sup>WF</sup>→RGC<sub>ON</sub> | 5 | -70.0 | 0.0 | 2.0 | -42.5 | 2.5 | I | 6.0 |
| BP<sub>OFF</sub>→RGC<sub>OFF</sub> | 13 | 0.0 | 0.0 | 2.5 | -44.0 | 3.0 | I | 6.0 |
| AMA<sub>OFF</sub><sup>WF</sup>→RGC<sub>OFF</sub> | 12 | -70.0 | 0.0 | 2.5 | -34.4 | 2.5 | I | 6.0 |
| AMA<sub>ON</sub><sup>NF</sup>→RGC<sub>OFF</sub> | 12 | -80.0 | 0.0 | 2.0 | -47.5 | 2.0 | I | 6.0 |

With the exception of RGCs, all cell types were modeled as leaky integrators, whose membrane potential (*v*_m_) followed the following differential equation:

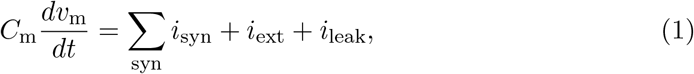

where *C*_m_ was the cell type–specific membrane conductance, the sum was over all presynaptic currents *i*_syn_ (see Section 4.2), *i*_ext_ was the external current resulting from extracellular electrical stimulation (see Section 4.5), and *i*_leak_ was a leakage current modeled by a constant linear conductance (*G*_m_) in series with a constant single-cell battery (i.e., the cell ‘s resting voltage, *E*_rest_):

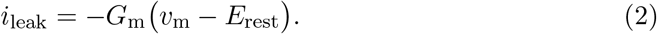

The light-cone interaction was modeled with an additional current as a synapse, described in detail in Cottaris and Elfar (2005) and given as:

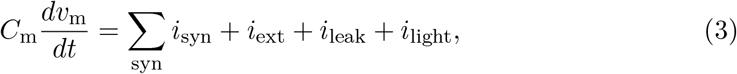

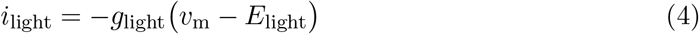

where *E*_light_ = − 8 mV was the reversal potential and *g*_light_ was the synaptic conductance that depended on the time-dependent light intensity

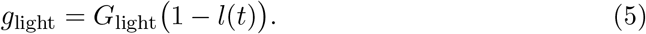

RGCs were modeled as Hodgkin-Huxley neurons with seven ion channels and a leakage current (*i*_leak_) to capture the spiking dynamics of the retina (Fohlmeister and Miller, 1997; Guo et al., 2016):

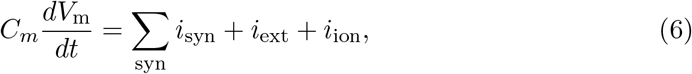

where the ionic current *i*_ion_ was given as the product of the neuron ‘s surface area *A* and the sum of several ionic current densities:

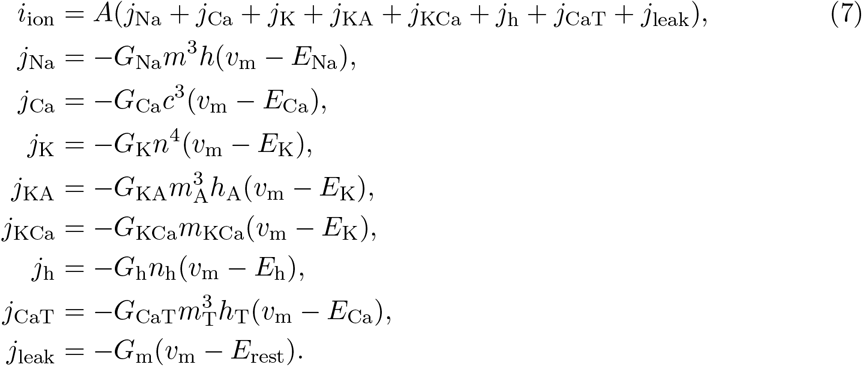

Here, *j*_Na_ was a voltage-gated sodium channel with gating variables *m* and *h*; *j*_Ca_ was a voltage-gated calcium channel with gating variable *c* and *E*_Ca_ modeled with the Nernst equation (see Guo et al. 2016 for details); *j*_K_ was a non-inactivating potassium channel with gating variable *n*; *j*_KA_ was an inactivating potassium channel with gating variables *m*_A_ (called *A* in Guo et al. 2016) and *h*_A_; *j*_KCa_ was a Ca^2+^-activated potassium channel gated by *m*_KCa_ whose value was dependent on the internal calcium concentration (see Guo et al. 2016 for details); *j*_h_ was a hyperpolarization-activated non-selective cationic channel with gating variable *n*_h_; and *j*_CaT_ was a low-threshold voltage-activated calcium channel with gating variables *m*_T_ and *h*_T_ (Fohlmeister and Miller, 1997; Guo et al., 2016).

The equations for the gating variables were identical to Guo et al. (2016) (see their Tables 2–3). In short, all gating variables, except the inactivating gating variable *h_T_* of *j_CaT_*, followed first-order kinetics:

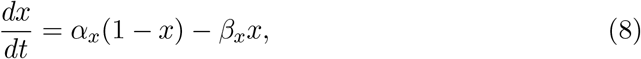

where *x* was the gating variable, *α_x_* was the opening rate and *β_x_* was the closing rate of the channel. The inactivating gating variable *h*_T_ of *j*_CaT_ (but not *m*_T_; note the typo in Guo et al. 2016) followed second-order dynamics:

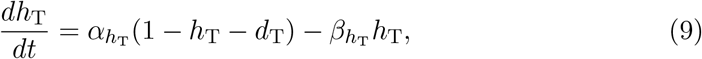

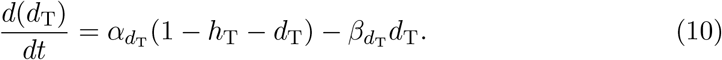

The initial values of the gating variables and the internal calcium concentrations of the RGCs can be found in Appendix A.4.

Neurons were assumed to contain a spherical soma with either 26 µm diameter in the case of RGCs (Milo et al., 2010; Crooks and Kolb, 1992) or 7 µm otherwise (Cottaris and Elfar, 2005). The initial values of the membrane voltages were set according to normal distributions, whose parameters can be found in Appendix A.4.

Ionic current densities were multiplied by the surface area *A* of the RGC to convert to a current. *G*_leak_ was set to a value so that the spontaneous firing rate under 0.5 light was around 2 Hz (Tao et al., 2020). A spike was recorded whenever the membrane potential exceeded − 10 mV.

### 4.2. Modeling the retinal circuitry

Inspired by Cottaris and Elfar (2005), we simulated a three-dimensional healthy patch (300 µ m × 300 µ m × 210 µ m) of the parafoveal retina. The network consisted of 11, 138 cells belonging to nine different cell types (4, 149 cones, 537 horizontal cells, 3, 508 ON/OFF bipolar cells, 779 ON/OFF wide-field amacrine cells, 723 narrow-field amacrine cells, and 1, 442 ganglion cells), connected via generally accepted (Rodieck, 1998; Wassle and Boycott, 1991) synaptic connections (see Fig. 1). For each type of neuron, the *x* and *y* coordinates were arranged hexagonally according to:

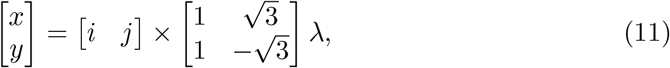

where *i, j* = 0, ± 1, ± 2… and *λ* was cell-type specific (Table 2). The *x* and *y* coordinates were further jittered by Gaussian noise: *N* (0, 1 µm). Neurons were confined to different *z* locations depending on their cell type (Table 2). Within each band, the *z* coordinate was assigned by sampling from a random uniform distribution.

Neurons were connected as shown in Fig. 1 using parameters given in Table 3. Synaptic delays (*τ* in Table 3) were set such that ON RGCs fired their first spikes roughly 20 ms after stimulus onset and OFF RGCs fired roughly 50 ms after stimulus onset (Zeck et al., 2011). To achieve these response times, we assumed a constant transmission speed and calculated the average distance between each connected pair of cells to set *τ* accordingly.

The synaptic connection from presynaptic neurons of the same type to a postsynaptic neuron was modeled with a current *i*_syn_ (see Equations 1, 3, and 6) in the postsynaptic neuron via:

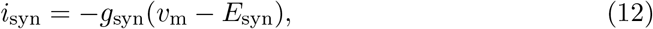

where *E*_syn_ was different for each synaptic type (see Table 3). *g*_syn_ was computed as a spatially weighted sum over the conductances of all the channels from the presynaptic neurons:

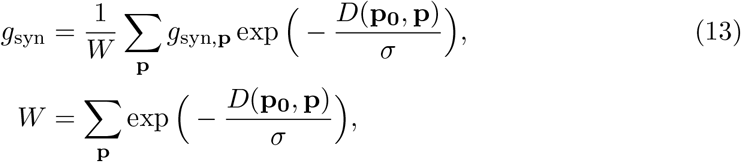

where *D*(**p_0_**, **p**) was the Euclidean distance between the center of the postsynaptic neuron **p_0_** and the center of a presynaptic neuron **p**, and *σ* was a decay constant that determined how the presynpatic currents were weighted. *σ* was different for each synaptic type (see Table 3). Following Cottaris and Elfar (2005), we modeled the individual channel conductance *g*_syn,**p**_ as either a monotonically increasing (type “I”) or monotonically decreasing (type “D”) function of the membrane potential of the presynaptic neuron with the following equation:

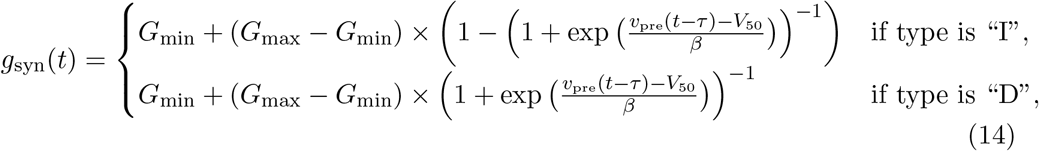

where *G*_min_ and *G*_max_ were the lower and upper bound of values for the synaptic conductance, *v*_pre_ was the membrane potential of the presynaptic neuron, *τ* was the synaptic delay, *V*_50_ determined the function ‘s center operating point and *β* determined the function ‘s steepness (see Table 3).

### 4.3. Modeling retinal degeneration

Retinal degenerative diseases are commonly described in the literature as progressing in three phases (Marc et al., 2003; Jones et al., 2016):

- Phase I starts with either cone or rod stress, which leads to the truncation of the outer segments. The population of the affected photoreceptors starts to decrease and their neurites start to extend.
- In Phase II, the other class of photoreceptors also start to die and extend their neurite. Cones continue to truncate. Muller cells move to the outer nuclear layer and start to seal off the retina from the choroid. This process is called subretinal fibrosis, which will later evolve to a glial seal. Horizontal cells begin to hypertrophy and extend their neurites, while rod and cone bipolar cells retract their dendrites.
- Phase III is a protracted period of cell death and leads to global retinal remodeling. In early Phase III, Muller cells hypertrophy and form the glial seal. Neurons start to die, while microneuromas start to form. They often contain active synapses despite lacking normal signaling abilities. In middle Phase III, progressive neuronal death and microneuroma formation continue, while the remaining neurons start to migrate. Specifically, amacrine and bipolar cells move to the inner plexiform and the ganglion cell layer, and amacrine cells and RGCs move to the glial seal. In late Phase III, the microneuromas regress with continued cell death, accompanied with the hypertrophy of Muller cells and vessels.

To make the modeling of such a complex process feasible, we limited ourselves to the major neuroanatomical changes that may have an impact on RGC signaling, and introduced them in a systematic step-wise manner. These changes are summarized in Table 4.

**Table 4.**
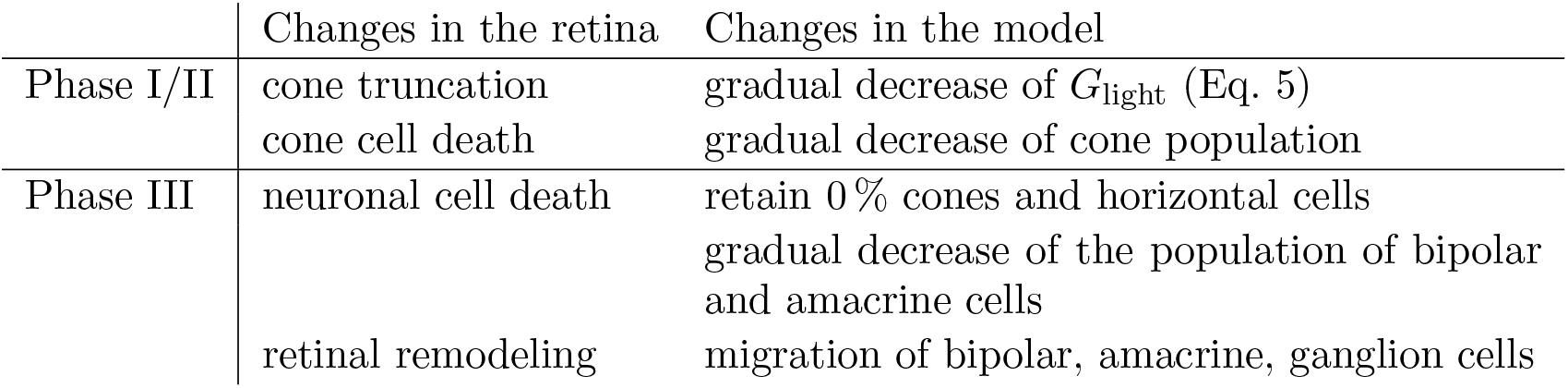
Phases of retinal degeneration (Marc et al., 2003), with corresponding changes in the retina and in our model.

|  | Changes in the retina | Changes in the model |
| --- | --- | --- |
| Phase I/II | cone truncation | gradual decrease of $G_{\text{light}}$ (Eq. 5) |
|  | cone cell death | gradual decrease of cone population |
| Phase III | neuronal cell death | retain 0 % cones and horizontal cells |
|  |  | gradual decrease of the population of bipolar and amacrine cells |
|  | retinal remodeling | migration of bipolar, amacrine, ganglion cells |

#### 4.3.1. Phase I/II

Because our model retina did not include rods, we combined Phases I and II into Phase I/II, where we gradually reduced the cone population and shortened the outer segments of the surviving cones, the latter of which was modeled by a gradual reduction in the ceiling of a cone ‘s light response, *G*_light_ (Eq. 5). To model disease progression over time (Fig. 4E), we assumed a linear reduction in cone segment length and cone population over time. The complete loss of photoreceptors and horizontal cells marked the end of Phase I/II.

#### 4.3.2. Phase III

To simulate global retinal remodeling in Phase III, we restricted ourselves to simulating cell death and migration.

First, we gradually reduced the population of bipolar and amacrine cells (but not RGCs). Second, according to the literature, a fraction of the surviving amacrine and bipolar cells tend to migrate to the inner plexiform layer and the ganglion cell layer, whereas amacrine cells and RGCs tend to migrate to the glial seal (Marc et al., 2003). We simulated this by migrating a randomly chosen subset of cells to different layers. A subset of amacrine cells was moved to the horizontal cell layer (*z ∈*[100 µm, 128 µm], close to the hypothetical glial seal), the inner plexiform layer (*z ∈*[40 µm, 80 µm]), and the ganglion cell layer (*z ∈*[25 µm, 39 µm]) in equal proportions. Half of the migrating bipolar cells were moved to the inner plexiform layer and the other half to the ganglion cell layer. The migrating RGCs were moved to the horizontal cell layer. The *z* coordinates of the migrating cells were sampled from a random uniform distribution in the respective range of *z* values that make up the different retinal layers (Table 5), whereas *x, y* coordinates remained unchanged. Synaptic weights and delays were unaffected by these coordinate changes.

**Table 5.** The range of *z* coordinates (*z*_min_, *z*_max_) where bipolars, amacrines and RGCs could be found during degeneration, given in µm. The bipolar cells migrated to the inner plexiform layer and the ganglion cell layer. The amacrine cells migrated to the horizontal cell layer, the inner plexiform layer and the ganglion cell layer. The RGCs migrated to the horizontal cell layer.

|  | BP<sub>ON/OFF</sub> | AMA<sub>ON/OFF</sub><sup>WF/NF</sup> | RGC<sub>ON/OFF</sub> |
| --- | --- | --- | --- |
| healthy ($z_{\min}, z_{\max}$) | (100, 128) | (80, 101) | (25, 39) |
| degenerated ($z_{\min}, z_{\max}$) | (25, 39) | (25, 39) | (100, 128) |
|  | (40, 80) | (40, 80) | - |
|  | - | (100, 128) | - |

To model disease progression over time (Fig. 4E), we assumed a linear reduction in cell survival rate (from 100 % at the beginning of Phase III to 0 at the end of Phase III) and a linear increase in cell migration rate (from 0 to 50 %).

Our model did not include Muller cells or microneuromas. However, some of the modeled changes may be indirectly due to Muller cell activity, such as the progressive death of inner retinal neurons.

### 4.4. Estimating spatiotemporal receptive fields

To measure the spatiotemporal receptive field of an RGC, we fit a generalized linear model (GLM) to its spiking response to a spatially correlated “cloud” stimulus (Shi et al., 2019). The stimulus consisted of spatiotemporal Gaussian white noise (pixel size = 6.25 µm, mean brightness = 0.5, standard deviation of brightness = 0.175, refresh rate = 20 Hz) filtered with a two-dimensional spatial Gaussian filter (standard deviation = 12.5 µm). The resulting spikes were binned at the refresh rate of the stimulus.

The generalized linear model predicted the firing rate of a RGC as:

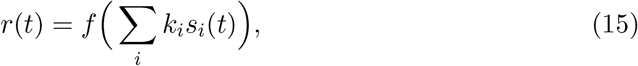

where *s_i_*(*t*) denoted the relevant stimulus frames before and at time *t, k_i_* denoted the spatial filter for the stimulus frame *s_i_*(*t*), and *f* was a nonlinear function. We trained a GLM in PyTorch with a spatiotemporal filter spanning five time steps and a rectified linear unit (ReLU) as the nonlinear function. The GLM was regularized with a Laplacian square penalty on the spatial filters (Shi et al., 2019). We used the mean squared error as loss function. The GLM was trained for 3000 epochs with the Adam optimizer and a learning rate of 0.000001 decayed by 10 % every 2000 epochs.

### 4.5. Modeling extracellular electrical stimulation

Extracellular electrical stimulation was assumed to be generated by a disk electrode with diameter *α* µm placed at (*x_e_, y_e_, z_e_*). The extracellular electrical potential *v*_ext_ at location (*x, y, z*) was given by:

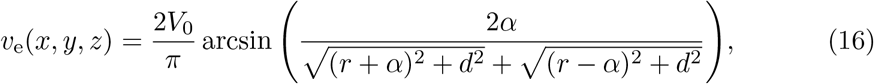

where *V*_0_ was the electrical potential of the disk,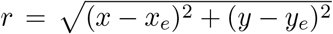
, and *d* = *z* − *z_e_* (Wiley and Webster, 1982). *v*_e_ was converted to a current, *i*_e_ (see Equations 1, 3, and 6), as follows:

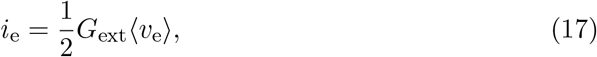

where *G*_ext_ was a conductance (see Table 2) and ⟨*v_e_*⟩ was the average of the absolute voltage differences between 500 uniformly sampled, diametrically opposing points on the neuron ‘s spherical soma (Knuth, 1969).

Although epiretinal electrodes are known to activate passing axon fibers (Rizzo et al., 2003a; Fried et al., 2009; Beyeler et al., 2019), we did not include RGC axons in our simulations.

The epiretinal electrode was centered at (*x, y, z*) = (0 µm, 0 µm, − 2 µm). The subretinal electrode was centered at (*x, y, z*) = (0 µm, 0 µm, 135 µm). Both electrodes had a diameter of *α* = 80 µm.

## Acknowledgements

This work was supported by the National Eye Institute of the National Institutes of Health under Award Number R00-EY029329. The content is solely the responsibility of the authors and does not necessarily represent the official views of the National Institutes of Health.

## Software and Data Availability

All our code will be made publicly available once the manuscript is accepted.

## Appendix

### A. Supplemental Information

#### A.1. Inter-spike intervals

Fig. A1A shows the inter-spike intervall (ISI) distribution for ON and OFF retinal ganglion cells (RGCs) as a function of disease progression. Fig. A1B highlights the histograms of the ISI distributions presented in Panel A during Phase I/II. Note that both the mean firing rate (see Fig. 4) and the mean ISI of ON cells (Fig. A1B, *left*) decreased as degeneration progressed. This is because most ON cells did not spike at all and therefore did not contribute any finite ISI values. However, a select few did spike a lot, therefore contributing many short ISI values. As a result, mean ISI decreased while mean firing rate decreased as well. Furthermore, we did not observe any bursting behavior in the spontaneous firing of RGCs.

**Figure A1.**
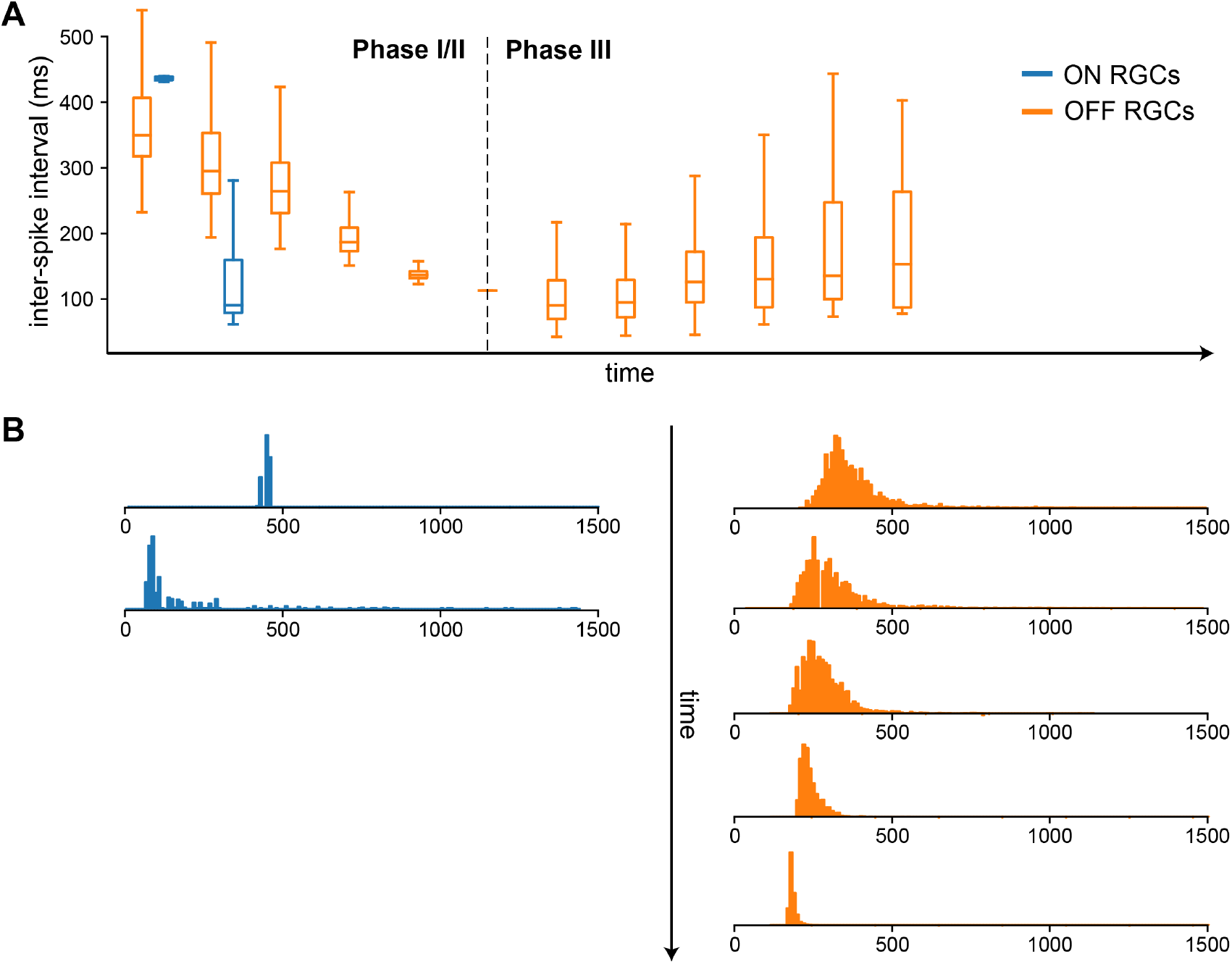
inter-spike intervall (ISI) distributions as a function of disease progression. **A)** Box plots of ISIs during Phase I/II and Phase III. **B)** Histograms for the data from Phase I/II that is presented in Panel A.

#### A.2. Phase I/II: Light response

Fig. A2 shows the firing rate of ON and OFF retinal ganglion cells (RGCs) in response to full-field stimuli of a given light intensity *l*(*t*) = const. (Eq. 5), averaged across the population of surviving cells, as a function of both cone outer segment truncation (simulated by a reduction in *G*_light_ and cone survival rate. As light intensity varied between 0 (black) and 1 (white), the RGC response to 0.5 intensity was interpreted

**Figure A2.**
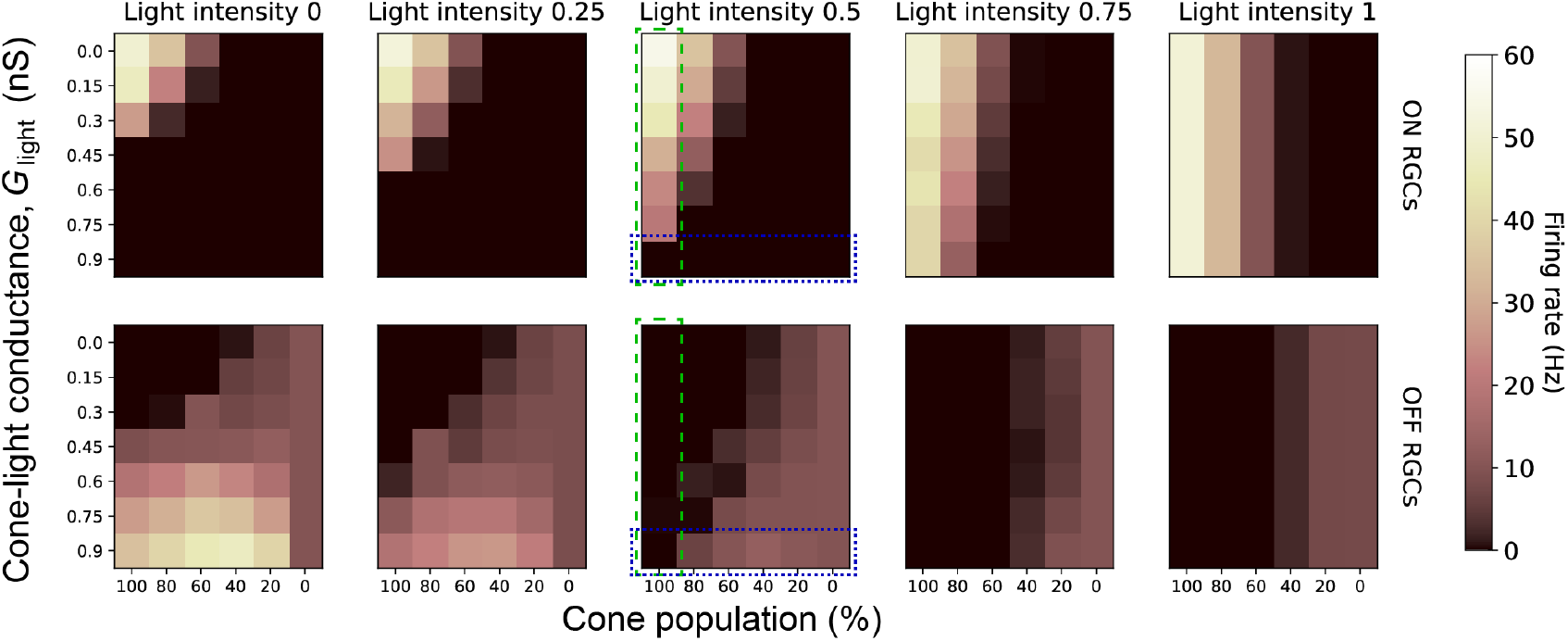
Light response of RGCs as a function of cone outer segment truncation and cone survival rate. The values encapsulated by the green dashed box are the same as the mean values presented in Fig. 4F. The values encapsulated by the blue dotted box are the same as the mean values presented in Fig. 4G.

At high light intensities (*l*(*t*) > 0.5), both ON and OFF RGC activity was mainly a function of the size of the surviving cone population. At low light intensities (*l*(*t*) ≤ 0.5), both cone outer segment truncation and cone population size had an effect on RGC firing, with ON firing dominated by outer segment truncation and OFF firing dominated by cone population. The linear time axis of disease progression corresponded to the diagonal in each heatmap (top-left corner: healthy, bottom-right corner: end of Phase I/II).

#### A.3. Ganglion cell response to electrical stimulation

Fig. A3 shows the RGC response to a constant electrical stimulus delivered either epiretinally (Fig. A3A) or subretinally (Fig. A3). The stimulus was the same as for the spatial profiles in Fig. 7; that is, a 20 Hz cathodic-first biphasic pulse train of 1 s duration, with pulse duration 0.45 ms and 60 µA amplitude. Electrode dimesnions are given in Section 4.5.

#### A.4. Initial values

Table A1 shows the initial values for the gating variables in the Hodgkin-Huxley model of the RGCs (Eq. 7). The gating variables are in units of Hz. The internal calcium concentration are in units of nM (nanomolar).

**Table A1.** Initial values for the Hodgkin-Huxley model.

| $m$ | $h$ | $c$ | $n$ | $A$ | $h_A$ | $y$ | $m_T$ | $h_T$ | $d_T$ | $[\text{Ca}^{2+}]_i$ |
| --- | --- | --- | --- | --- | --- | --- | --- | --- | --- | --- |
| 0.5 | 0.5 | 0.5 | 0.5 | 0 | 1 | 0.5 | 0 | 1 | 1 | 0.0001 |

Table A2 shows the initial values of the membrane potential for the different cell types. Initial values for each cell type followed a normal distribution with a mean *µ* and a standard deviation *σ*.

**Table A2.** Initial values for the membrane potential of the different cell types.

| Cell type | $\mu$ (mV) | $\sigma$ (mV) |
| --- | --- | --- |
| cone photoreceptors | -46.8 | 3 |
| horizontal cells | -47.7 | 3 |
| ON bipolar cells | -35 | 3 |
| OFF bipolar cells | -44 | 3 |
| ON wide-field amacrine cells | -42 | 3 |
| OFF wide-field amacrine cells | -34.5 | 3 |
| ON narrow-field amacrine cells | -47.6 | 3 |
| ON RGCs | -66.5 | 3 |
| OFF RGCs | -70.5 | 3 |

**Figure A3.**
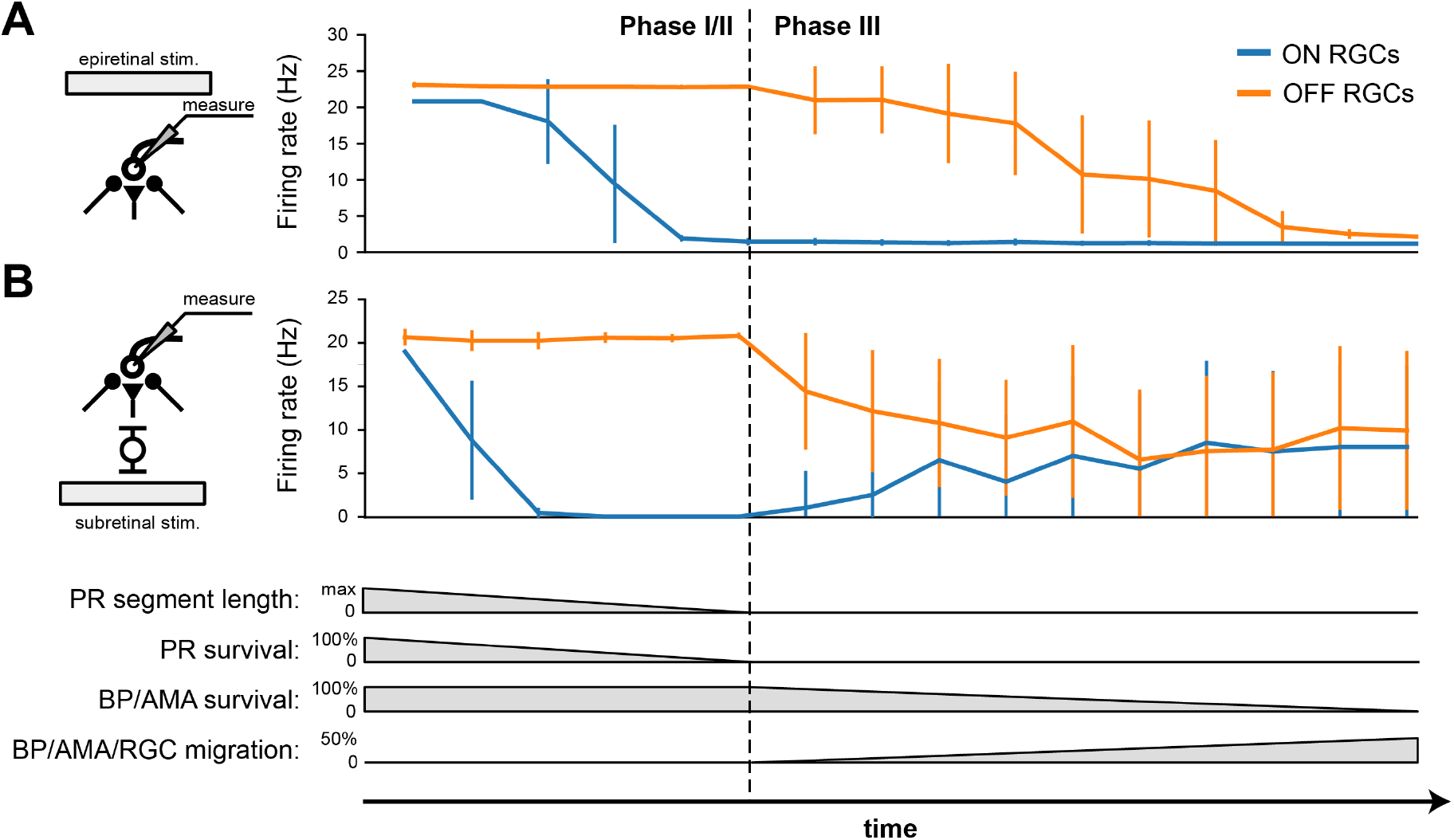
RGC firing rate in response to epiretinal (A) and subretinal (B) stimulation. The mean and standard deviation were calculated from the ON and OFF RGCs that were located directly under the electrode.

## Notes

### Competing Interest Statement

The authors have declared no competing interest.

